# Kiawah and Seabrook islands are a critical site for the *rufa* Red Knot (*Calidris canutus rufa*)

**DOI:** 10.1101/2022.03.21.485188

**Authors:** Mary Margaret Pelton, Sara R. Padula, Julian Garcia-Walther, Mark Andrews, Robert Mercer, Ron Porter, Felicia Sanders, Janet Thibault, Nathan R. Senner, Jennifer A. Linscott

**Author notes:** Corresponding authors &. These authors contributed equally (Mary Margaret Pelton, Sara R. Padula).

## Abstract

The *rufa* Red Knot (*Calidris canutus rufa*) is a migratory shorebird that performs one of the longest known migrations of any bird species — from their breeding grounds in the Canadian Arctic to their nonbreeding grounds as far south as Tierra del Fuego — and has experienced a population decline of over 85% in recent decades. During migration, knots rest and refuel at stopover sites along the Atlantic Coast, including Kiawah and Seabrook islands in South Carolina. Here, we document the importance of Kiawah and Seabrook islands for knots by providing population and stopover estimates during their spring migration. We conducted on-the-ground surveys between 19 February - 20 May 2021 to record the occurrence of individually marked knots. In addition, we quantified the ratio of marked to unmarked knots and deployed geolocators on knots captured in the area. Using a superpopulation model, we estimated a minimum passage population of 17,247 knots (~41% of the total *rufa* knot population) and an average stopover duration of 47 days. Our geolocator results also showed that knots using Kiawah and Seabrook islands can bypass Delaware Bay and fly directly to the Canadian Arctic. Finally, our geolocators, combined with resighting data from across the Atlantic Flyway, indicate that a large network of more than 70 coastal sites mostly concentrated along the coasts of Florida, Georgia, South Carolina, and North Carolina provide stopover and overwintering habitat for the knots we observed on Kiawah and Seabrook islands. These findings corroborate that Kiawah and Seabrook islands should be recognized as critical sites in the knot network and, therefore, a conservation priority. As a result, the threats facing the sites — such as prey management issues, anthropogenic disturbance, and sea level rise — require immediate attention.

## INTRODUCTION

Migration is the process whereby individuals move from one area to another to exploit alternative resources or environments that fluctuate in suitability (Winger *et al.* 2019). Migration can range from short-distance movements to those that cover tens of thousands of kilometers (Alerstam *et al.* 2003). The *rufa* Red Knot (*Calidris canutus rufa,* hereafter, ‘knot’) is a long-distance migratory shorebird that performs one of the longest migrations of any bird species (Piersma *et al.* 2005, Conklin *et al.* 2017), with some individuals traveling ~30,000 km from their breeding grounds in the Canadian Arctic (70°N) to nonbreeding grounds in the southeastern U.S. (Niles *et al.* 2012) and as far south as Tierra del Fuego at the southern tip of South America (53 – 54°S; Niles *et al.* 2008, Burger *et al.* 2012). Knots face numerous threats along their migratory route and, in 2015, were listed as a threatened species in both Canada and the United States due to a steep population decline of 85% over the past few decades (USFWS 2014).

Knots migrate between their breeding and nonbreeding grounds along the Atlantic Flyway and undertake several nonstop flights of thousands of kilometers without feeding or resting (Niles *et al.* 2008). To endure the energetic demands of such long flights, knots rely on stopover sites where they can rest and refuel (Baker *et al.* 2004, Atkinson *et al.* 2007). The annual cycles and migratory strategies of knots, in turn, require that knots time their migrations to coincide with the occurrence of superabundant but ephemeral resources at their stopovers (Piersma & Baker 2000). Delaware Bay — one of the best studied stopover sites for knots — exemplifies this dependency. Knot arrival at Delaware Bay coincides with the spawning of the horseshoe crabs (*Limulus polyphemus)* on which they feed in order to refuel and continue their migrations to the Arctic (Niles *et al.* 2008, McGowan *et al.* 2011, Tucker *et al.* 2021). Delaware Bay, however, represents only one site within a larger network of beaches, estuaries, and barrier islands (Cohen *et al.* 2009, 2010a,b, Tuma & Powell 2021) that share a suite of pressures including anthropogenic disturbance (Burger *et al.* 2007), coastal development (Buler & Moore, 2011), and unsustainable prey harvest practices (Niles *et al.* 2009). As a result, knots’ foraging efficiency has decreased at some sites, and they have failed to reach the minimum threshold of mass gain to complete their migrations in some years (Baker *et al.* 2004). Given that climate change and sea level rise will continue to exacerbate the pressures on stopover sites that support knots and other shorebirds (Iwamura *et al.* 2013, Rakhimberdiev *et al.* 2018), there is an urgent need to identify all critical stopover sites for these at-risk species.

Within North America, the network of sites used by knots spans much of the Atlantic Coast. In addition to Delaware Bay, the Eastern Shore of Virginia is known to support upwards of 5,000 knots during spring migration (Cohen *et al.* 2010a). Recent evidence suggests that more than two dozen coastal sites in the southeastern U.S. (i.e., from Texas to South Carolina) also support knots during migration and the nonbreeding season (Tuma & Powell 2021). In Georgia, for instance, estimates indicate that between 8,000-24,000 knots stop during fall migration (Lyons *et al.* 2018). Intriguingly, flocks of up to 8,000 knots have been recorded in spring as well on the Kiawah-Seabrook Island complex in South Carolina (Thibault 2013) — a relatively small site in comparison to most others in the network. Kiawah and Seabrook islands, however, have received relatively little attention and much remains to be learned about how (and how many) knots use them over the course of the year (Smith *et al.* 2019).

Population estimates of overwintering and migrating shorebirds are traditionally difficult to quantify because of fluctuating numbers of birds as they enter and exit a site during the season (Lyons *et al.* 2016, Lok *et al.* 2019). We aim to build on the count data collected on Kiawah and Seabrook islands in previous studies to assess its importance to knots by accounting for the flow-through nature of stopover sites. To do this, we used a mark-resighting approach following Lyons *et al.* (2016) to estimate the: (1) population size, (2) stopover duration, (3) connectivity, and (4) overwintering status of knots using the Kiawah-Seabrook Island complex. We complemented these analyses by assessing stopover site usage along the Atlantic Flyway by knots carrying light-level geolocators and using the online http://www.bandedbirds.org database to link the knots we resighted in South Carolina with resightings from other sites across the flyway. Ultimately, we believe that our study can contribute to a broader understanding of the knot annual cycle and efforts to conserve knots wherever they occur throughout the year.

## METHODS

### Study area

Our study area comprised 24 kilometers of sandy beach in the Kiawah-Seabrook Island complex (hereafter, KSI) on the coast of South Carolina (32°35′10.6“N, 80°07′47.6“W; Fig. 1). Seabrook and Kiawah islands are bordered by the North Edisto and Stono rivers, respectively. The islands are divided by Captain Sam’s Inlet, which is critical to local fauna, including strand feeding common bottlenose dolphins (*Tursiops truncates*; Dybas 2021), foraging seabirds, and a variety of roosting shorebirds, including knots. This area also provides critical overwintering habitat for Piping Plovers (*Charadrius melodus*) and nesting habitat for Wilson’s Plovers (*C. wilsonia*), American Oystercatchers (*Haematopus palliatus*), and Least Terns (*Sternula antillarum*), all of which are species of conservation concern in South Carolina (USFWS 2001, SCDNR 2015). Due to its unusual hydrodynamic regime, the inlet experiences frequent shifts and geomorphological changes. It has also been intentionally relocated several times, most recently in 2015 (Doyle & Adams 2015). A semi-diurnal tide results in shorebirds using the inlet twice each day to roost during high tide.

**Figure 1.**
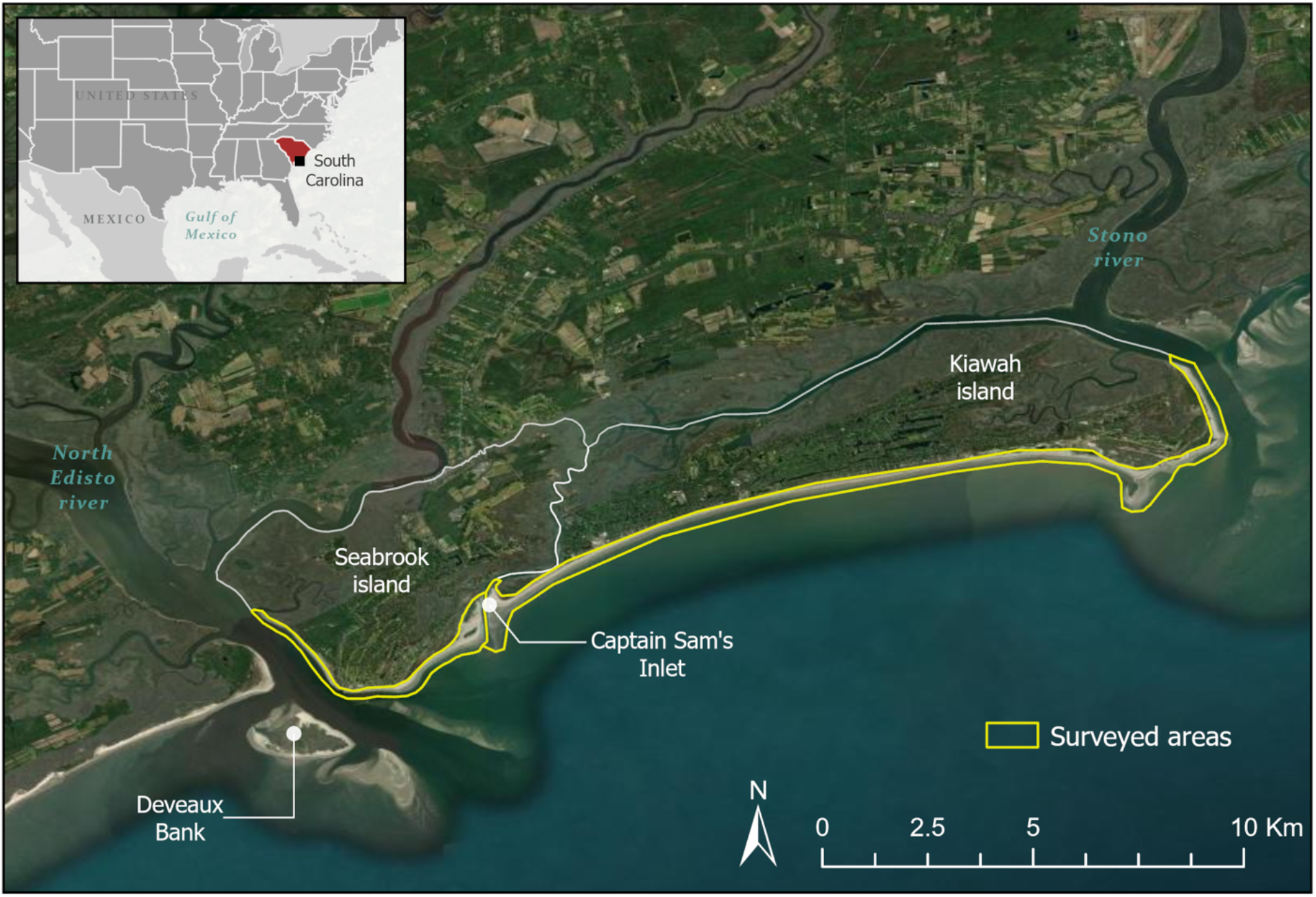
The Kiawah-Seabrook Island Complex, South Carolina consists of 24 kilometers of sandy beaches and an estuarine inlet.

At low tides, shorebirds occupy the complex’s sandy beaches where they feed primarily on invertebrate prey. Benthic core and knot fecal samples suggest the knots diet on KSI consists primarily of coquina clams (*Donax variabilis*; Thibault & Levisen 2013). Both islands are popular tourist destinations with fully developed coastlines containing golf courses, several resorts, and popular public beach access points with high levels of human and dog disturbance. At the south end of the study area, South Carolina Department of Natural Resources owned Deveaux Bank — an ephemeral offshore sand bank — provides additional undeveloped habitat for shorebirds and marine birds. Knots move back and forth between KSI and Deveaux Bank, especially during spring horseshoe crab spawning events, with horseshoe crab eggs comprising an additional important component of their diet (Thibault & Levisen 2013, SCDNR 2018). As Deveaux Bank requires boat access, it was not surveyed during our study.

### Banding

Standardized shorebird banding was implemented in the Western Hemisphere beginning in the mid-1980s (PASG 2016). Across the Atlantic Flyway, ~8% (Lyons 2021) of the knot population is marked with flags consisting of a unique alphanumeric code that allows for the identification of each marked bird (Clark *et al.* 2005). Flags are colored according to the Pan American Shorebird Group protocol, with colors corresponding to the area in which the bird was marked (PASG 2016). These possible areas are: Canada, the United States, Mexico, Central America, the Caribbean, northern South America, Argentina, and Chile. By resighting knots that have been previously marked by other researchers, we estimated their passage population and stopover duration. Compiling resighting data from other sites in North America also allowed us to identify connections between South Carolina and stopover sites throughout the knot migratory network.

### Data collection

We collected data from 19 February - 20 May 2021 — a period during which migrating knots were likely to be present in KSI. After a two-week period of observer training, KSI was surveyed 2-4 times a week by two separate observers between February-March. In April and May — the period of expected peak knot presence — we increased our sampling effort to 4-6 times a week. These observations resulted in 13 weeks of total effort. Due to restricted access, flag readings and scan samples were collected from Seabrook, but not Kiawah, from 14-20 May.

The 24 km of KSI beaches were surveyed by randomly selecting beach-walking access points, which served as the survey starting point for a given island. We performed surveys within ~2.5 hours of high tide when knots were likely to be relatively stationary. When a flock was found, we performed a series of flag ratio scans using spotting scopes by recording the number of individuals with and without leg flags. We sampled without replacement as much as possible by scanning from one end of the flock to the other. Efforts were also made to avoid recounting flocks in the same day by not taking samples from flocks that we had already encountered and from which we therefore recognized flag combinations. Additionally, we ensured that we had adequate coverage of birds on KSI to the best of our ability by randomly selecting sites to survey and having two observers working separately during the week. If birds in a scan had flags, we recorded the flag color, alphanumeric code, and presence of any other device that it might carry. We also recorded the specific location, time, wind, and tide of each sighting.

### Flag and geolocator deployment

Between April 2015 and May 2016, we deployed geolocators on knots captured in South Carolina. Knots were captured using cannon nets on Bird Key Stono Seabird Sanctuary on 21 April 2015; in Cape Romain National Wildlife Refuge at Marsh Island on 16 October 2015; and on Deveaux Bank Seabird Sanctuary on 10 May 2016. Captures were made at high tide roosts at Bird Key and Marsh Island, and while feeding on horseshoe crab eggs as the tide fell on Deveaux Bank. Upon capture, birds were immediately removed from the net and placed in keeping cages for processing. All birds were measured and fitted with a uniquely inscribed 3-character leg flag and U.S. Geological Survey metal band. Migrate Technology Ltd. geolocators weighing 1.1 g were attached to leg bands as described in Niles *et al.* (2010). The units record the maximum light level every minute and conductivity (i.e., contact with alkaline water) every 3 seconds. A total of 33 geolocators were deployed, but only three were retrieved. One knot captured at Deveaux Bank was recaptured on Tierra del Fuego on 13 January 2018. A geolocator deployed at Bird Key was retrieved from a dead knot (presumably from red tide, a harmful algal bloom) found in the Tampa Bay, Florida region. And, finally, a geolocator from Marsh Island was retrieved from a knot recaptured at Seabrook Island on 29 April 2017.

### Mark-resighting analysis

We reconstructed mark-resighting data for each marked knot on a weekly basis by combining all observations within one week into a single observation, beginning 19 February. We used the http://www.bandedbirds.org database to verify our resightings. For those marking schemes that had been input into the database (e.g., from the United States and Canada), we removed observations of flag codes that had no corresponding capture history. Some marking schemes did not appear in the database, however; in these cases, we retained all observations with a high degree of observer certainty. We then used an open population Jolly-Seber (JS) modeling framework to explore the flow of arriving, stopped over, and departing knots during the spring migratory period. Specifically, we used the superpopulation parameterization of the JS model — a hierarchical, state-space parameterization that incorporates data augmentation (Crosbie & Manly 1985, Royle & Dorazio 2008, Kéry & Schaub 2012) — to estimate the probabilities of entering (β) the study site, staying (ϕ) to the next week, and resighting (ρ). These parameters are analogous to the demographic parameters of recruitment, survival, and recapture in conventional mark-recapture JS models, respectively. We implemented methods from Lyons *et al.* (2016) to add a binomial model for flag ratio scan samples, which also enabled us to estimate total passage population, or the number of knots estimated to use KSI during the spring migratory period. More recently developed models additionally allow for the statistical identification of groups that might pass through a site separately (e.g., Lok *et al.* 2019). However, knot local- and regional-scale movements, and the resulting resighting heterogeneity early during our study period (see below), precluded us from employing these models, meaning that we could only estimate a single, average stopover duration for all knots using KSI.

We verified that the JS model fit our data using goodness-of-fit tests developed for capture-recapture models in the R Programming Environment (version 4.1.1; R Core Development Team 2016) and package ‘R2ucare’ (Gimenez *et al.* 2018). Omnibus testing of the null hypothesis that the model was an adequate fit for the data was non-significant (*p* = 0.11, χ^2^ = 43.34, df = 33), indicating that we did not need to adjust the model for transience or trap-dependence. We also explored support for time-varying versus constant parameters. In a stopover context, entry probabilities (β) are commonly allowed to vary over time, since individuals may be less likely to arrive at a stopover site during some weeks (e.g., the last week) than others. We compared various models that constrained the probabilities of staying (ϕ) and resighting (ρ) to be constant and/or allowed them to vary with sampling occasion (week). Models were fit using encounter histories and the POPAN model in program MARK (White and Burnham 1999) *via* the R package ‘RMark’ and compared using Akaike’s Information Criterion adjusted for small sample sizes (AIC_c_; Burnham & Anderson 2002). The most parsimonious model (AIC_c_< 2 and with the fewest parameters) consisted of a constant staying probability and weekly variation in resighting probability (Table 1).

**Table 1.** Model comparison for time-varying versus constant ɸ and p parameters. K = number of parameters; Weight = Akaike weight.

| Model | K | AIC<sub>c</sub> | ΔAIC<sub>c</sub> | Weight | Deviance |
| --- | --- | --- | --- | --- | --- |
| $\beta_t \phi_c \rho_t$ | 27 | 856.594 | 0.000 | 0.999 | -711.943 |
| $\beta_t \phi_t \rho_t$ | 38 | 879.294 | 22.700 | <0.001 | -716.126 |
| $\beta_t \phi_c \rho_c$ | 15 | 897.980 | 46.385 | <0.001 | -643.268 |
| $\beta_t \phi_t \rho_c$ | 26 | 905.934 | 49.339 | <0.001 | -660.251 |

We performed a Bayesian analysis integrating the superpopulation JS model with the binomial scan sample model, using the approach outlined in Lyons *et al.* (2018). The JS model estimates stopover duration using encounter histories and a latent state variable that reflects arrival in and departure from the study site (Lyons *et al.* 2016). The latent state variable (*z*_i,t_) is a time-specific Bernoulli random variable for each individual *i* in the population (i.e., flagged and unflagged) at time *t*, where *z*_i,t_= 1 while using the stopover site and *z*_i,t_= 0 before arrival and after departure. The posterior latent states are then summed across individuals to estimate the stopover duration (Lyons *et al.* 2018). In this way, the calculation of mean stopover duration is robust to variations in resighting probability, which were pronounced during weeks when knots roosted and foraged on Deveaux Bank or moved among sites in the southeastern U.S.

Using flag scan samples, we modeled the number of flagged individuals in a scan sample as a binomial random variable

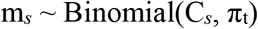

where m_*s*_ is the number of flagged individuals in the scan sample *s*, C_*s*_ is the number of individuals checked for flags, and π_t_ is the proportion of flagged individuals in the population during the corresponding week. This modification for weekly variation in the proportion of the population that is flagged (π_t_) suits the pulsed arrivals of two or more migratory groups that may differ in the proportion flagged, which we expected to find at our study site. The total passage population was then calculated as the sum of the arrivals at each sampling occasion, adjusted for the flagged proportion at the corresponding occasion

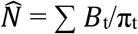

where 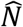 is the total passage population, *B*_t_, is the number of entries/arrivals at a sampling occasion *t*, and π_t_ is the proportion of the population carrying alphanumeric flags at occasion *t.* In this way, the integrated model thus estimated both stopover dynamics (e.g., stopover duration, mean staying probability, weekly resighting probability) and the derived weekly and total passage population.

The full model incorporating the JS model and flag scan sample ratios was implemented in R using the package ‘jagsUI’ (Kellner 2016) as an interface with JAGS software (Plummer 2003). We followed guidance from Kéry & Schaub (2012) for parameter-expanded data augmentation (PX-DA; Royle & Dorazio 2012) in order to fix the parameter space for analysis, adding potential unobserved individuals as all-zero encounter histories (*n* = 600) to our dataset. We checked that this value was sufficiently large by visually inspecting the posterior distribution of the estimated number of total marked individuals (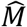) to verify that it was not truncated. Following Lyons *et al.* (2018), we used uniform priors (0,1) for the staying and resighting probabilities and uninformative priors for arrival probability. We simulated three MCMC chains of 90,000 iterations each, with burn-in periods of 30,000 iterations and a thinning rate of 3. We assessed convergence of the chains visually and *via* the Brooks-Gelman-Rubin statistic (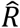; Brooks & Gelman 1998), where models with parameter values < 1.1 were considered to have converged. All values are presented as means and 95% credible intervals unless otherwise noted.

### Geolocator and connectivity analysis

To corroborate our mark-resighting results, we used our geolocation data to answer three questions: (1) How long did knots stopover in South Carolina during their northward migration? (2) How many stops did knots make after departing from South Carolina for the breeding grounds? and, (3) Where were those stops? Geolocators were analyzed using the R package *FLightR* (Rakhimberdiev *et al.* 2017) — which has been shown to successfully delineate the movements of migratory shorebirds (Rakhimberdiev *et al.* 2016) — and followed the workflow outlined in Lisovski *et al.* (2019). Briefly, before deploying our geolocators, we placed them outside for a ~7-day long period that we then used within *FLightR* to calibrate the data from each geolocator. We otherwise used the default package settings, but constrained the locations identified as stopover sites to coastal areas. We included all sites at which an individual was estimated to stop for at least 2 days with a (movement) probability of 0.8 and considered an individual to have stopped in South Carolina if a stopover was recorded between 30-33°N. Using this approach, we estimated the mean (± 1 SD) of the dates of arrival and departure from South Carolina, the number of stops an individual made after departing South Carolina on its way northward, and the location of those stops.

To further assess migratory connectivity, we searched the online database http://www.bandedbirds.org and retrieved the capture and resighting history for every knot that we resighted during our study. With this information, we mapped the network of sites used by knots during their passage through the U.S. and Canada. We eliminated sites that were less than 10 km from each other using a rarifying filter in R (Brown *et al.* 2014).

## RESULTS

### Resighting results

We recorded 217 uniquely flagged knots during our 13-week study period. Resightings occurred in all weeks except for the week beginning 26 March, when knots in the KSI area were observed only on nearby Deveaux Bank (N.R. Senner pers. obs.). We recorded flag scan samples in all weeks except for the two-week period beginning 19 March, also due to temporary knot roosting and foraging on Deveaux Bank. Resightings and scan samples occurred throughout KSI but were most common near Captain Sam’s Inlet, shortly before or after high tide. Most resighted individuals carried dark green or lime green flags indicative of having been flagged in the U.S. Other individuals flagged in South America appeared later in the season (Fig. 2) and during this period we observed flocks of at least 4,000 individuals on multiple occasions. While new resightings occurred up until the final week, nearly all knots had departed by 21-27 May (M. Andrews & R. Mercer, pers. obs.).

**Figure 2:**
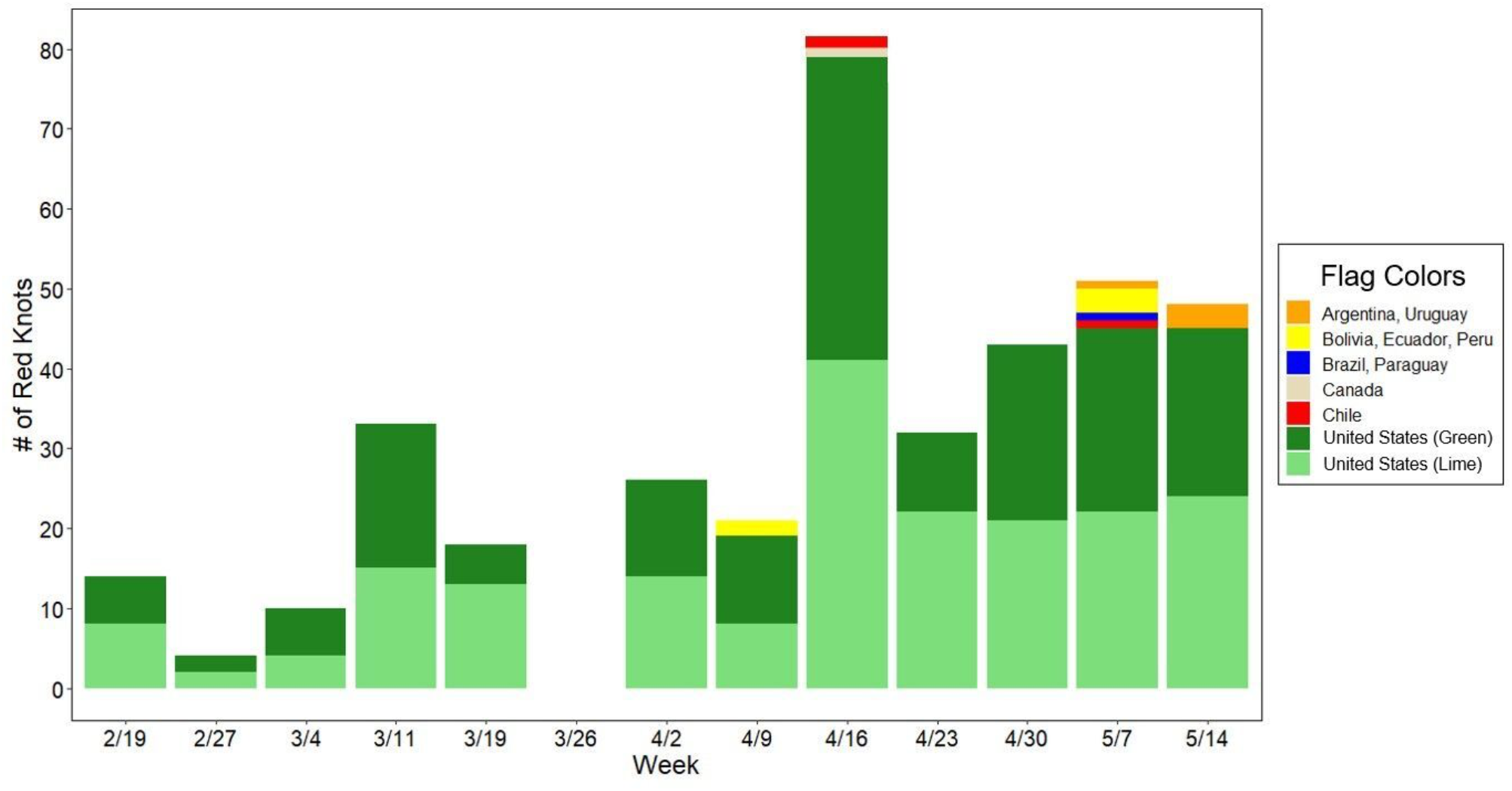
Number of uniquely flagged Red Knots and the corresponding flag colors encountered each week from February-May 2021 on Kiawah and Seabrook islands, South Carolina. Dates shown represent the start of each week.

Our model estimated a minimum total passage population of 17,247 knots (95% CI: 13,548, 22,099) during our study period. Of this passage population, 2.4% (95% CI: 1.9, 2.9) on average were estimated to be flagged. This estimate is approximately 41% of the total global population of 42,000 *rufa* knots (Andres *et al.* 2012). On average, individuals spent 47 days (95% CI: 40.1, 54.8) at the study site.

The model estimated one constant parameter — staying probability (0.91, 95% CI: 0.86, 0.95) — and three parameters that were allowed to vary on a weekly basis — entry probability, resighting probability, and population size. Entry probability was highest in the first week and remained relatively consistent thereafter (Fig. 3A). Resighting probability increased in the fourth week, was lower during the weeks that knots spent most of their time on Deveaux Bank, and increased again later in the season (Fig. 3B). The estimated weekly population size increased throughout the season, starting at 2,411 (95% CI: 858, 5,050) in the first week and ending at 13,852 (95% CI: 9,764, 19,254) the week of 14 May (Fig. 3C).

**Figure 3.**
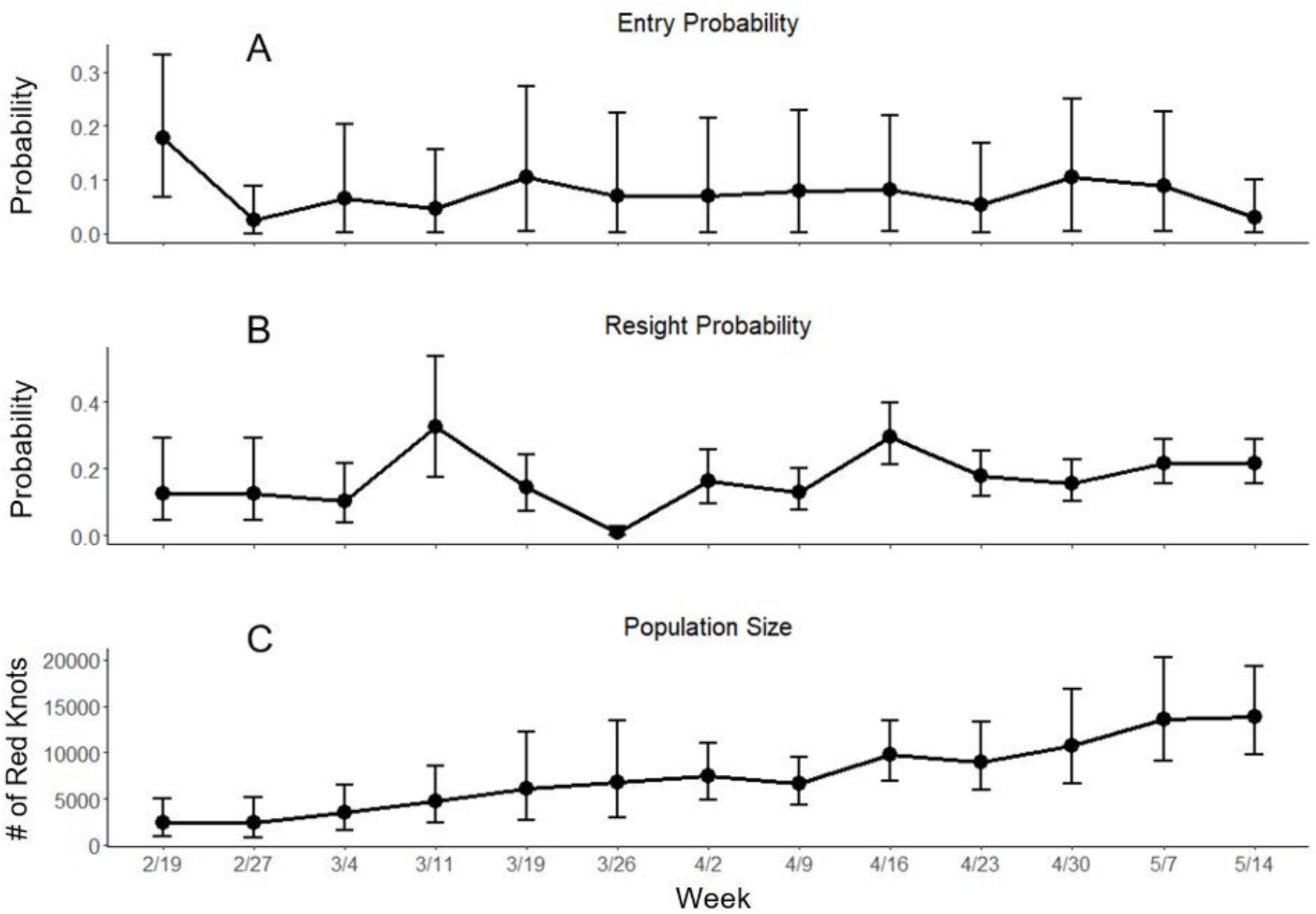
Model-derived estimates of the weekly entry probability (A), resighting probability (B), and population size (C) for Red Knots on Kiawah and Seabrook islands, South Carolina. Bars around estimates represent model generated 95% credible intervals.

### Geolocator results

Each of our three geolocator-carrying individuals spent the nonbreeding season in a different region: one on Tierra del Fuego (50.54°S; Fig. 4A, B); one along the Atlantic Coast, moving among sites from Georgia to North Carolina (30.3-35.1°N; Fig. 4C); and one on the Gulf Coast of Florida (27.2-29.2°N; Fig. 4D). Of the two individuals that did not spend the nonbreeding season in or close to South Carolina, both arrived in the region 2-5 May and departed approximately three weeks later *(μ* = 20 ± 1 d), between 23 May-1 June (*n* = 3 departures). Across all three individuals and all years during which they were tracked, northward departure from South Carolina averaged 24 May ± 5 d (*n* = 5 departures). Once departing South Carolina, the three individuals stopped on average 1.8 ± 1 times, with those stops occurring north of 49°N, mostly along the western shore of either James or Hudson bays, Canada (49.3-55.4°N; Fig. 4A). Because of the nature of geolocation data and Arctic summers, we were unable to identify the breeding areas used by any of the individuals.

**Figure 4:**
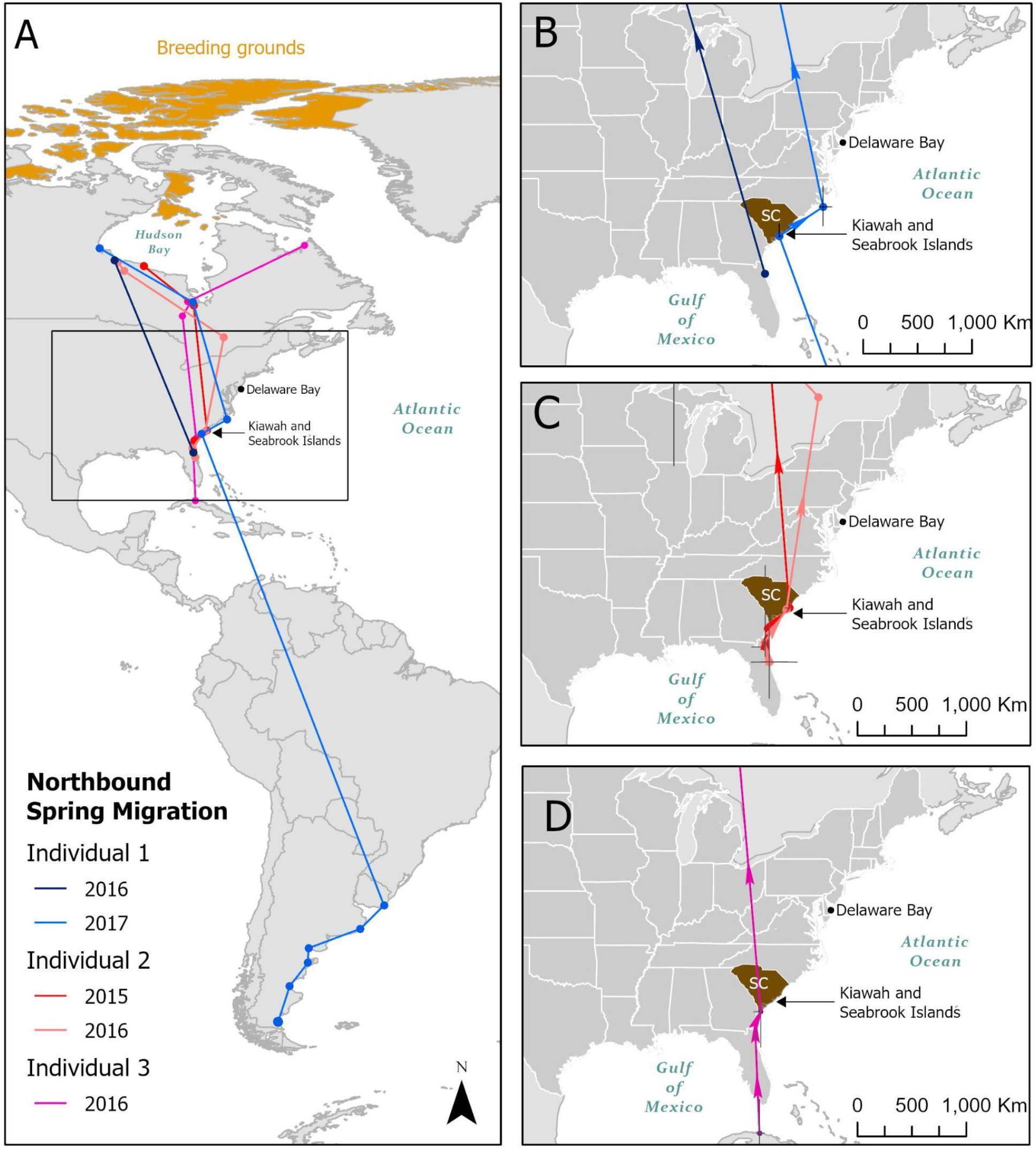
Northbound migratory tracks obtained using light-level geolocators attached to three Red Knots (blue, red, and purple lines, respectively) from 2015-2017 (map inset A). Map inset B, C, and D show northbound movements of individuals 1, 2, and 3 along the Atlantic Coast of the United States, respectively. Vertices show latitudinal and longitudinal 95% confidence intervals (black lines). Kiawah and Seabrook islands are highlighted to show when birds stopped over at the site, while Delaware Bay is shown as a reference point for other studies of knot migration.

### Flyway-wide resighting results

The capture history of the birds with dark or lime green flags that we resighted indicates that they were captured at 18 sites along the U.S. Atlantic Coast (1 in Texas, 4 in Florida, 4 in South Carolina, and 8 in Delaware Bay). Furthermore, based on their resighting history, the knots we resighted have used a network of at least 74 sites, including beaches, barrier islands, estuaries, inlets, and sandbars. These resightings came from Texas, Louisiana, Florida, Georgia, South Carolina, North Carolina, Virginia, Maryland, Delaware, New Jersey, Connecticut, and Massachusetts in the U.S., as well as Ontario (James Bay) and Quebec in Canada.

## DISCUSSION

The Atlantic Coast of North America hosts a number of critical migratory stopover sites for *rufa* Red Knots, but the majority of scientific and conservation attention has focused on only a few of those sites, such as Delaware Bay (Baker *et al.* 2004, Atkinson *et al.* 2007, Niles *et al.* 2009) and coastal Virginia (Cohen *et al.* 2009, 2010a,b). Using a superpopulation model, we estimated that the 24-km stretch of sandy beaches on Kiawah and Seabrook islands in South Carolina hosted at least 17,247 (95% CI: 13,548, 22,099) knots from February-May 2021, representing ~41% of the estimated global *rufa* knot population of 42,000 individuals (Andres *et al.* 2012). While knot site fidelity to individual sites may exhibit large interannual variation (Piersma *et al.* 2021, Tucker *et al.* 2021) depending on the conditions and resource dynamics of the region (van Gils *et al.* 2005), our study suggests that KSI is a critical site for knots along the Atlantic Coast and deserves increased recognition and conservation attention.

### Population estimates

Across the Atlantic Flyway, a combination of methods has been used to develop estimates of knot population sizes at stopover and nonbreeding sites. These have ranged from: (1) mark-recapture approaches, which led to an estimation of 18,000 knots using Delaware Bay during spring migration in 2004 (Gillings *et al.* 2010) and 23,400 knots using the Georgia coast in fall 2011 (Lyons *et al.* 2018); (2) peak count approaches, which resulted in estimates of 5,939 knots using the Virginia coast from 2004-2007 (Cohen *et al.* 2010a); and, (3) hybrid approaches, which generated estimates of 8,750 knots using four sites along the South Carolina and Georgia coasts in the spring of 2019 (Smith *et al.* 2019). This variation in methodologies — as well as variation in their interpretation — has led to persistent uncertainty about the number of knots using specific sites, as well as about the total number occurring along the Atlantic Flyway. We hypothesize that our higher estimates of knots using KSI relative to previous estimates from the site and those from other sites along the Atlantic Coast may be a result of either interannual variation in site usage as suggested by Tucker *et al.* (2021) and our geolocator data (Fig. 4), improved resighting effort and statistical methodologies, an actual shift in site usage by knots, or a combination of the three. Our estimates should nonetheless be viewed as a minimum, as additional knots might have passed through KSI the week of 21-27 May after our regular resighting efforts ceased. Regardless, this variation corroborates the need for more comprehensive estimates to be generated from mark-recapture studies along the entire Atlantic Coast on a regular basis.

### KSI likely supports both overwintering and migrant knots

Our resighting (Fig. 1) and geolocator (Fig. 4) data indicate that there are likely two groups of knots using KSI: (1) overwintering knots that stayed on KSI and in surrounding areas or arrived early in the year, as exemplified by the abundance and substantial number of flags we detected at the beginning of our surveys, and (2) spring migrants, as exemplified by the increased population estimates and proportion of flagged individuals we observed as the season progressed. Knots are known to use the southeastern U.S. coast, including South Carolina, during the nonbreeding season (Burger *et al.* 2012, Niles *et al.* 2012, Tuma & Powell 2021), and Lyons *et al.* (2018) estimated that this region alone supports ≥ 10,400 knots during this period. Our results confirmed that one of our geolocator-carrying individuals moved among Georgia, South Carolina, and North Carolina during the nonbreeding season before ultimately migrating northward from South Carolina in late May. Likewise, during our surveys, we estimated that ~2,400 knots were present at KSI as early as 19 February (Fig. 2). We then subsequently detected ~42% of the marked knots we observed during the first two weeks of the study again in May. In contrast, we would expect that longer distance migrants would exhibit later arrivals and shorter stopover durations. Accordingly, our other two geolocator-carrying knots — which spent the nonbreeding season in Florida and southern South America, respectively (Fig. 4) — arrived in South Carolina in early May and departed ~3 weeks later. This suggests that KSI may not only be an important stopover site for spring migrants but an important site for overwintering knots as well.

In this respect, our results mirror those of Lyons *et al.* (2018), who found that during fall migration knots spent an average of 38 days at the Altamaha River Delta, with a group of longer distance migrants (Tierra del Fuego) that stopped over for ~21 days and a group of shorter distance migrants (Southeastern U.S., Caribbean islands, and Brazil) that stopped over for ~42 days. Our model, however, was unable to differentiate between the two apparent groups in our study and we could not estimate the difference in the duration of their respective stays on KSI. This is because the methods from Lyons *et al.* (2016), on which our models were based, are unable to identify the presence of multiple groups and our resighting effort was not great enough to parameterize more complex models (e.g., Lok *et al.* 2019) that could capture the varying behaviors likely exhibited by the ‘overwintering’ and ‘migrant’ groups. A key focus of future research should therefore be to increase resighting efforts in the region, and specifically on KSI, throughout the nonbreeding season (early November – early June) to try to generate robust estimates of the sizes of these two apparent groups.

### Importance of KSI in the knot migratory network

Delaware Bay has historically been regarded as the last steppingstone at which knots can refuel before departing to their Arctic breeding grounds (Baker *et al.* 2004). Much conservation attention has thus been focused on the site (Clark 1993, Atkinson *et al.* 2007, Niles *et al.* 2009, McGowan *et al.* 2011). However, an increasing body of evidence suggests that knots rely on a suite of stopover sites across the flyway for a variety of purposes (Cohen *et al.* 2010a, Tuma & Powell 2021). Our survey of the http://www.bandedbirds.org database to connect the knots we resighted on KSI with other sites across the Atlantic Flyway corroborates these recent studies and revealed a network of more than 70 sites spanning Texas to Maryland (Fig. 5). Our geolocator results also indicate that Delaware Bay is likely not the only terminal stopover site used by knots prior to reaching the Canadian Arctic: the five spring migration departures we obtained from our three geolocator-carrying individuals indicate that they all skipped Delaware Bay after stopping in South Carolina (Fig. 4). These geolocator results are supported by the results of a recent study that used an automated radio telemetry array to track knots from South Carolina on their northward migration (Smith *et al. in prep*.) and which found that the majority of these knots skipped Delaware Bay and went directly to the Arctic from South Carolina. Nonetheless, resighting data from http://www.bandedbirds.org also suggested that a substantial number of the knots stopping over in South Carolina are subsequently resighted in Delaware Bay in at least some years. Further tracking work, combined with on-the-ground efforts to quantify knot refueling rates and social dynamics, would therefore help clarify the migratory stopover decisions that result in different stopover behaviors being exhibited from year to year (e.g., Chan *et al.* 2019, Linscott & Senner 2021).

**Figure 5.**
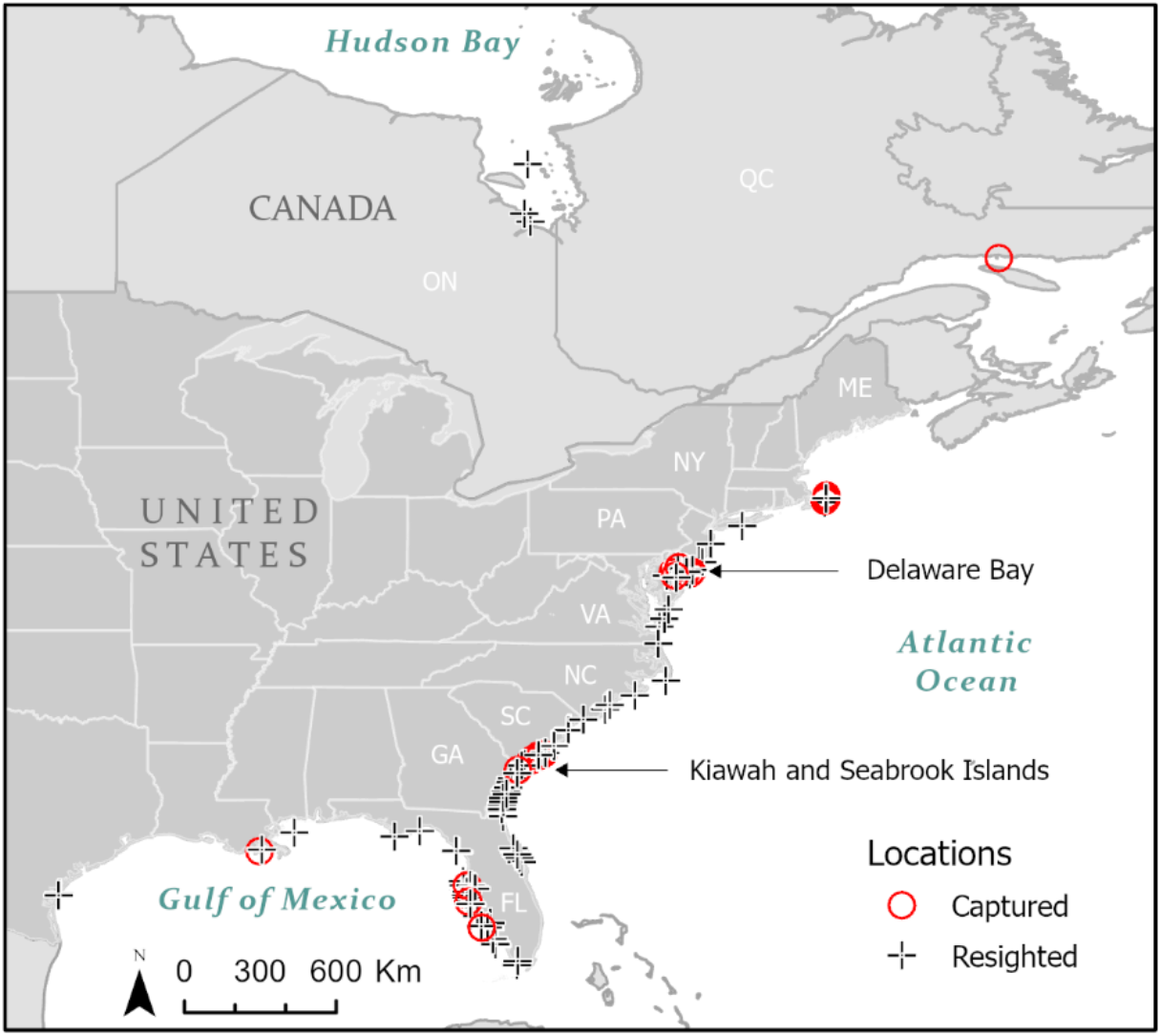
Network of sites where the Red Knots resighted on Kiawah and Seabrook islands, South Carolina were initially flagged (red circles) and where they were subsequently resighted (black crosses) along the Atlantic Coast of Canada and the U.S.

Regardless of whether knots are regularly using KSI and the greater South Carolina coast as a terminal stopover site, the comparable importance of KSI as a migratory stopover site for knots is clear. Though KSI is relatively small, we estimate that 41% of the total knot population is passing through during migration. What is more, a nocturnal roost supporting nearly half of the Atlantic Flyway population of Hudsonian Whimbrel (*Numenius hudsonicus*) was recently discovered on Deveaux Bank (Sanders *et al.* 2021), a site we also observed knots using during our study. Within the framework of the Western Hemisphere Shorebird Reserve Network, KSI (and Deveaux Bank) would therefore qualify as a site of Hemispheric Importance (WHSRN 2021). The fact that KSI only comprises 24 km of beaches — with Deveaux Bank comprising just a few more — underscores its importance and, likely, sensitivity to conservation threats such as those associated with human disturbances, habitat degradation, and outright habitat loss.

Results from the loss and degradation of stopover sites used by knots elsewhere in the Atlantic Flyway (Baker *et al.* 2004) and across the globe (Studds *et al.* 2017) point to the consequences for the knot migratory network that such changes could cause (Xu *et al.* 2019). The importance of KSI is amplified even further if knots are indeed not only migrating through KSI but also overwintering there. In such a scenario, knots would be reliant on the site for most of their nonbreeding season (~6 months) and not just the spring migration period (~2-3 months).

### Conservation implications

Because of the variability of knot migratory patterns and their on-going population declines (Piersma *et al.* 2021, Tucker *et al.* 2021), it is critical to recognize the threats that knots are facing across the Atlantic Flyway. Delaware Bay, for instance, has suffered a > 75% decline in its use by knots since 1990, which Niles *et al.* (2009) suggested is likely related to declines in horseshoe crab numbers. To refuel sufficiently, knots need healthy populations of their prey, such as horseshoe crab eggs. At terminal stopover sites like Delaware Bay from which knots leave for their breeding grounds, they must be able to refuel and meet a certain weight threshold (e.g., > 180 g) to successfully complete their migrations (McGowan *et al.* 2011). Female crab abundance at Delaware Bay has thus been shown to positively correlate with an individual’s ability to reach this body mass threshold and survive to subsequent years (Baker *et al.* 2004). Recent results from the Cape Romain National Wildlife Refuge in South Carolina indicate a similar positive correlation between horseshoe crab spawning and knot densities at the refuge (Takahashi *et al.* 2021). In our study area, knots appear to primarily feed on coquina clams on KSI itself, as horseshoe crabs do not currently occur on the islands, but horseshoe crabs do spawn on Deveaux Bank and knots are known to feed on their eggs there (Thibault & Levinsen 2013). Maintaining and understanding the diversity of knot prey used in South Carolina — including both horseshoe crabs and coquina clams — is therefore important in order to enable knots to rapidly refuel and successfully complete their northward migrations.

Anthropogenic disturbance also poses a threat to knots that must focus their time and energy on foraging and building up sufficient pre-departure fat reserves to continue their northward migrations (Thomas *et al.* 2003). Burger *et al.* (2007) found that knots and other shorebirds using sites with little human disturbance were able to spend 70% of their time foraging, while their foraging efficiency was reduced by more than 40% at high disturbance sites. Because KSI is highly developed, with tourist attractions such as golf courses, public beach access points, and resorts, there is substantial human and dog traffic along its beaches. During our 13-week survey period we observed that, with the onset of spring, increasing numbers of people and dogs led to the frequent disturbance of knots and other shorebirds. Currently both islands do have dog leash laws intended to decrease the disturbance of wildlife: some areas are off limits to dogs year-round, while in others, dogs are allowed either on and/or off leash, depending on the time of day and year. For large portions of the two islands, however, these latter restrictions do not cover the entirety of the period that knots are likely present. On Kiawah Island, dogs are allowed off leash across much of the island from 1 November - 15 March, while on Seabrook Island, dogs can be off leash in the central portion of the island during the evening and early morning (17:00-09:59 hrs), even from 1 April - 30 September (Towns of Kiawah Island and Seabrook Island). Our results suggest that these rules should be extended and enforced throughout the period knots are present on the islands.

Climate change poses an additional variety of unpredictable challenges to knots and their associated coastal habitats. The Mid-Atlantic Coast of the U.S. has exhibited some of the fastest rates of sea level rise in the world (∼4-10 mm y^−1^; Ezer & Corlett 2012). Because much of the Mid-Atlantic Coast (including KSI) is highly developed, increased inundation may cause the loss of critical habitat by squeezing beaches and mudflats in between existing infrastructure (von Holle *et al.* 2019). Sea level rise may also alter the geomorphology and hydrodynamic regimes of estuaries (Khojasteh *et al.* 2021), potentially altering key roosting sites such as Captain Sam’s Inlet and Deveaux Bank. Subsequently, a reduction in habitat as a result of sea level rise (and other anthropogenic factors) could further increase human disturbance, as well as inter- and intraspecific competition for roosting and foraging areas (Goss-Custard 1988), thereby constraining the ability of knots to refuel (Baker *et al.* 2004). Efforts to preserve the integrity of the full suite of habitats currently found on KSI in the face of these potential changes, such as through beach replenishment and habitat migration strategies, is a major priority.

### Conclusions

We provide evidence of the critical importance of the Kiawah-Seabrook Island complex as a stopover and overwintering site for knots. In order to preserve the site’s ability to sustain shorebird populations, increased recognition and protection of the site is a priority. To this end, we recommend the nomination of KSI and neighboring Deveaux Bank as a Site of Hemispheric Importance in the Western Hemisphere Shorebird Reserve Network, as our results show that more than 40% of the *rufa* knot population, as well as 50% of the Atlantic Flyway Hudsonian Whimbrel population (Sanders *et al.* 2021), use the area. Because KSI may be serving as a terminal stopover site for knots, conservation measures should also focus on maintaining adequate abundances of knot prey to enable them to sufficiently refuel for nonstop flights to the Arctic. Human and dog disturbance on the islands lead to decreased foraging efficiency and use of roosting sites (Koch & Paton 2013); our study’s estimates of knot stopover duration on KSI can be used to help inform the timing of leash laws and other restrictions. Finally, sea level rise is expected to be a source of difficulty for knots in terms of the loss of suitable foraging and roosting habitat, and should remain a factor in management decisions given the critical importance of Captain Sam’s Inlet and the adjacent KSI beaches.

## ACKNOWLEDGEMENTS

We are very grateful for the help of the Senner Lab at the University of South Carolina, biologists at the South Carolina Department of Natural Resources, and the volunteers from the Kiawah Island Shorebird Stewardship Program and the Seabrook Island Birders. We also thank L. Usyk from http://www.bandedbirds.org for providing data and assistance, as well as Y. Aubry, H. Bellman, J. Brush, N. Martínez Curci, K. Kalasz, S. Koch, D. Newstead, L. Niles, and R. Rodrigues for permission to use their data. Helpful comments on early drafts were provided by J. Fraser, J. Lyons, S. Karpanty, T. Piersma, M. Stager, and M. Verhoeven. This research was supported by the South Carolina Department of Natural Resources and federal funding from South Carolina State Wildlife Grants to FS and JT, as well as a grant from the Magellan Scholars Program at the University of South Carolina to MP and SP, and a Carolina Bird Club Research Grant to SP. Finally, thanks to L. Niles for advice on geolocator deployment and retrieval, and M. Chaplin with the U.S. Fish and Wildlife Service for support

## LITERATURE CITED

Alerstam, T., A. Hedenström & S. Akesson. 2003. Long-distance migration: evolution and determinants. Oikos 103: 247–260.

Andres, B.A., P.A. Smith, R.I.G. Morrison, C.L. Gratto-Trevor, S.C. Brown & C.A. Friis. 2012. Population estimates of North American shorebirds. Wader Study Group Bulletin 119: 178–194.

Atkinson, P.W., A.J. Baker, K.A. Bennett, N.A. Clark, J.A. Clark, K.B. Cole, A. Dekinga, A. Dey, S. Gillings, P.M. González, K. Kalasz, C.D.T. Minton, J. Newton, L.J. Niles, T. Piersma, R.A. Robinson & H.P. Sitters. 2007. Rates of mass gain and energy deposition in Red Knot on their final spring staging site is both time‐ and condition‐dependent. Journal of Applied Ecology 44: 885–895.

Banded Birds. 2022. Shorebird Resighting Database: Banding Data Extraction: http://www.bandedbirds.org. Retrieved March 6, 2022.

Baker, A.J., P.M. Gonzalez, T. Piersma, L.J. Niles, I. de Lima Serrano do Nascimento, P.W. Atkinson, N.A. Clark, C.D.T. Minton, M.K. Peck & G. Aarts. 2004. Rapid population decline in Red Knots: fitness consequences of refuelling rates and late arrival in Delaware Bay. Proceedings of the Royal Society of London B 271: 875–882.

Berkson, J. & C.N. Shuster Jr., 1999. The Horseshoe Crab: the battle for a true multiple-use resource. Fisheries 24: 6–10.

Brooks, S.P. & A. Gelman. 1998. General methods for monitoring convergence of iterative simulations. Journal of Computational and Graphical Statistics 7: 434–455.

Brown, J. L. 2014. SDMtoolbox: a python-based GIS toolkit for landscape genetic, biogeographic and species distribution model analyses. Methods in Ecology and Evolution 5: 694–700.

Buler, J.J., & F.R. Moore. 2011. Migrant–habitat relationships during stopover along an ecological barrier: extrinsic constraints and conservation implications. Journal of Ornithology 152: 101–112.

Burger, J., S.A. Carlucci, C.W. Jeitner & L.J. Niles. 2007. Habitat choice, disturbance, and management of foraging shorebirds and gulls at a migratory stopover. Journal of Coastal Research 23: 1159–1166.

Burger, J., L.J. Niles, R.R. Porter, A.D. Dey, S. Koch & C. Gordon. 2012. Migration and over-wintering of Red Knots (*Calidris canutus rufa*) along the Atlantic Coast of the United States. Condor 114: 302–313.

Burnham, K.P. & D. R. Anderson. 2002. Model selection and multimodel inference: a practical information-theoretic approach. Springer, New York, New York, USA.

Chan, Y.-C., T.L. Tibbitts, T. Lok, C.J. Hassell, H.-B. Peng, Z. Ma, Z.W. Zhang & T. Piersma. 2019. Filling knowledge gaps in a threatened shorebird flyway through satellite tracking. Journal of Applied Ecology 56: 2305–2315.

Clark, K.E., L.J. Niles & J. Burger. 1993. Abundance and distribution of migrant shorebirds in Delaware Bay. Condor 95: 694–705.

Clark, N.A., S. Gillings, A.J. Baker, P.M. González & R. Porter. 2005. The production and use of permanently inscribed leg flags for waders. Wader Study Group Bulletin 108: 38–41.

Cohen, J., S.M. Karpanty J.D. Fraser, B.D. Watts & B.R. Truitt. 2009. Residence probability and population size of Red Knots during spring stopover in the Mid-Atlantic region of the United States. Journal of Wildlife Management 73: 939–945.

Cohen, J.B., S.M. Karpanty & J.D. Fraser. 2010. Habitat selection and behavior of Red Knots on the New Jersey Atlantic Coast during spring stopover. Condor 112: 655–662.

Cohen, J.B., S.M. Karpanty, J.D. Fraser & B.R. Truitt. 2010. The effect of benthic prey abundance and size on Red Knot (*Calidris canutus*) distribution at an alternative migratory stopover site on the US Atlantic Coast. Journal of Ornithology 151: 355–364.

Conklin, J.R., N.R. Senner, P.F. Battley & T. Piersma. 2017. Extreme migration and the individual quality spectrum. Journal of Avian Biology 48: 19–36.

Crosbie, S.F. & B.F.J. Manly. 1985. Parsimonious modelling of capture-mark-recapture studies. Biometrics 41:385–398.

Doyle, B.C. & M.R. Adams. 2015. Statistical evaluation of shoreline change: A case study from Seabrook Island, South Carolina. Environmental & Engineering Geoscience 21: 165–180.

Dybas, C.L. 2021. High-stakes mudbank chase: At low tide, US Southeast dolphins “beach” their prey. Oceanography 34: https://doi.org/10.5670/oceanog.2021.404.

Ezer, T. & B. Corlett. 2012. Is sea level rise accelerating in the Chesapeake Bay? A demonstration of a novel approach for analyzing sea level data. Geophysical Research Letters. 39: L19605.

Gimenez, O., E. Cam & J.M. Gaillard. 2018. Individual heterogeneity and capture‐recapture models: What, why and how? Oikos 127: 664–686.

Gillings S., P.W. Atkinson, A.J. Baker, K.A. Bennett, N.A. Clark, K.B. Cole, P.M. González, K.S. Kalasz, C.D.T. Minton, L.J. Niles, R.C. Porter, I.D.L. Serrano, H.P. Sitters & J.L. Woods. 2009. Staging behavior in Red Knot (*Calidris canutus*) in Delaware Bay: implications for monitoring mass and population size. Auk 126: 54–63.

Goss-Custard, J.D., J.T. Cayford & S.E.G. Lea. 1998. The changing trade-off between food finding and food stealing in juvenile oystercatchers. Animal Behavior 55: 745–760.

Iwamura T., H.P. Possingham, I. Chadès, C. Minton, N.J. Murray, D.L. Rogers, E.A. Treml & R.A. Fuller. 2013. Migratory connectivity magnifies the consequences of habitat loss from sea-level rise for shorebird populations Proceedings of the Royal Society B 280: 20130325.

Kellner, K. 2016. jagsUI: a wrapper around “rjags” to streamline “JAGS” analyses v. 1.4.4. R Foundation for Statistical Computing, Vienna, Austria. Accessed 20 February 2022.

Kéry, M. & M. Schaub. 2012. Bayesian population analysis using WinBUGS: a hierarchical perspective, 1st ed. Academic Press, Waltham, Massachusetts.

Khojasteh, D., W. Glamore, V. Heimhuber & S. Felder. 2021. Sea level rise impacts on estuarine dynamics: a review. Science of The Total Environment 780: 146470.

Koch, S.L. & P.W. Paton. 2014. Assessing anthropogenic disturbances to develop buffer zones for shorebirds using a stopover site. Journal of Wildlife Management 78: 58–67.

Lisovski, S., S. Bauer, M. Briedis, S.C. Davidson, K.L. Dhanhal-Adams, M.T. Hallworth, J. Karagicheva, C.M. Meier, B. Merkel, J. Ouwehand, L. Pederson, E. Rakhimberdiev, A. Roberto-Charron, N.E. Seavy, M.D. Sumner, C.M. Taylor, S.J. Wotherspoon & E.S. Bridge. 2020. Light-level geolocator analyses: a user’s guide. Journal of Animal Ecology 89: 221– 236.

Linscott, J.A. & N.R. Senner. 2021. Beyond refueling: investigating the diversity of functions of migratory stopover events. Ornithological Applications 123: 1–14.

Lok, T., C.J. Hassell, T. Piersma, R. Pradel & O. Gimenez. 2019. Accounting for heterogeneity when estimating stopover duration, timing and population size of Red Knots along the Luannan Coast of Bohai Bay, China. Ecology and Evolution 9: 6176–6188.

Lyons J., W. Kendall, J. Royle, S. Converse, B. Andres & J. Buchanan. 2016. Population size and stopover duration estimation using mark-resight data and Bayesian analysis of a superpopulation model. Biometrics 72: 262–271.

Lyons, J.E., B. Winn, T. Keyes & K.S. Kalasz. 2018. Post‐breeding migration and connectivity of Red Knots in the western Atlantic. Journal of Wildlife Management 82: 383–396.

Lyons, J.E. 2021. Red Knot stopover population size and migration ecology at Delaware Bay, USA, 2021. A report submitted to the Adaptive Resource Management Subcommittee and Delaware Bay Ecosystem Technical Committee of the Atlantic States Marine Fisheries Commission.

McGowan, C.P., J.E. Hines, J.D. Nichols, J.E. Lyons, D.R. Smith, K.S. Kalasz, L.J. Niles, A.D. Dey, N.A. Clark, P.W. Atkinson, C.D.T. Minton & W. Kendall. (2011). Demographic consequences of migratory stopover: linking Red Knot survival to horseshoe crab spawning abundance. Ecosphere 2: 69.

Niles, L.J., H.P. Sitters, A.D Dey, P.W. Atkinson, A.J. Baker, K.A. Bennett, R. Carmona, K.E. Clark, N.A. Clark, C. Espoz, P.M. González, B.A. Harrington, D.E. Hernández, K.S. Kalasz, R.G. Lathrop, R.N. Matus, C.D.T. Minton, R.I.G. Morrison, M.K. Peck, W. Pitts, R.A. Robinson & I.L. Serrano. 2008. Status of the Red Knot (*Calidris canutus rufa*) in the Western Hemisphere. Studies in Avian Biology 36: 1–36.

Niles, L.J., J. Bart, H.P. Sitters, A.D. Dey, K.E. Clark, P.W. Atkinson, A.J. Baker, K.A. Bennett, K.S. Kalasz, N.A Clark, K.J. Clark, S. Gillings, A.S. Gates, P.M. González, D.E. Hernández, C.D.T. Minton, R.I.G. Morrison, R.R. Porter, R.K. Ross & C.R. Veitch. 2009. Effects of horseshoe crab harvest in Delaware Bay on Red Knots: are harvest restrictions working? BioScience 59: 153–164.

Niles, L., J. Burger, R.R. Porter, A. Dey, C.D.T. Minton, P.M. Gonzalez, A.J. Baker, J.W. Fox & C. Gordon. 2010. First results using light level geolocators to track Red Knots in the Western Hemisphere show rapid and long intercontinental flights and new details of migration paths. Wader Study Group Bulletin 117: 1–8.

Niles, L.J., J. Burger, R.R. Porter, A.D. Dey, S. Koch, B. Harrington, K. Iaquinto & M. Boarman. 2012. Migration pathways, migration speeds and non-breeding areas used by northern hemisphere wintering Red Knots *Calidris canutus* of the subspecies *rufa*. Wader Study Group Bulletin 119: 195–203.

Pan American Shorebird Group. 2016. Pan American Shorebird Program Shorebird Marking Protocol. https://www.shorebirdplan.org/wp-content/uploads/2016/08/PASP-Marking-Protocol-April-2016.pdf.

Piersma, T. & A.J. Baker. 2000. Life history characteristics and the conservation of migratory shorebirds. In L.M. Gosling & W.J. Sutherland (Eds.), Behaviour and conservation (pp. 105–124). Cambridge: Cambridge University Press.

Piersma, T., D.I. Rogers, P.M. Gonzalez, L. Zwarts, L.J. Niles, I. S. de Lima do Nascimento, C.D.T. Minton & A.J. Baker. 2005. Fuel storage rates before northward flights in Red Knots worldwide: facing the severest ecological constraint in tropical intertidal environments? In: Greenberg, R. and Marra, P. P. (eds.), Birds of two worlds: ecology and evolution of migration. Baltimore: Johns Hopkins University Press, pp. 262–273.

Piersma, T., E.M.A. Kok, C.J. Hassell, H.-B. Peng, Y.I. Verkuil, G. Lei, J. Karagicheva, E. Rakhimberdiev, P.W. Howey, T.L. Tibbitts & Y.-C. Chan. 2021. When a typical jumper skips: itineraries and staging habitats used by Red Knots (*Calidris canutus piersmai*) migrating between northwest Australia and the New Siberian Islands. Ibis 163: 4, 1235–1251.

Plummer, M. 2003. JAGS: Just Another Gibbs Sampler. https://sourceforge.net/projects/mcmc-jags/. Accessed 20 February 2022.

Rakhimberdiev, E., N.R. Senner, M.A. Verhoeven, D.W. Winkler, W. Bouten & T. Piersma. 2016. Comparing inferences of solar geolocation data against high-precision GPS data: annual movements of a double-tagged Black-tailed Godwit. Journal of Avian Biology 47: 589–596.

Rakhimberdiev, E., A. Saveliev, T. Piersma & J. Karagicheva. 2017. FLightR: an r package for reconstructing animal paths from solar geolocation loggers. Methods in Ecology and Evolution 8: 1482–1487.

Rakhimberdiev, E., S. Duijns, J. Karagicheva, C.J. Camphuysen, V.R.S. Castricum, A. Dekinga, R. Dekker, A. Gavrilov, J. ten Horn, J. Jukema, A. Saveliev, M. Soloviev, T.L. Tibbitts, J.A. van Gils & T. Piersma. 2018. Food abundance at refuelling sites can mitigate Arctic warming effects on a migratory bird. Nature Communications 9, 4263.

Rehfisch, M.M., G.E. Austin, S.N. Freeman, M.J.S. Armitage & N.H.K. Burton. 2004. The possible impact of climate change on the future distributions and numbers of waders on Britain’s non-estuarine coast. Ibis 146: 70–80.

Royle, J.A. & R.M. Dorazio. 2008. Hierarchical modeling and inference in ecology: the analysis of data from populations, metapopulations and communities. Academic Press, London, UK.

Royle, J.A. & R.M. Dorazio. 2012. Parameter-expanded data augmentation for Bayesian analysis of capture-recapture models. Journal of Ornithology 152: S521–S537.

Sanders, F.J., M.C. Handmaker, A.S. Johnson & N.R. Senner. 2021. Nocturnal roost on South Carolina coast supports nearly half of the Atlantic coast population of Hudsonian Whimbrel Numenius hudsonicus during northward migration. Wader Study 128: 117–124.

Shimazaki, H., M. Tamura, Y. Darman, V. Andronov, M.P. Parilov, M. Nagendran & H. Higuchi. 2004. Network analysis of potential migration routes for Oriental White Storks (*Ciconia boyciana*). Ecological Research 19: 683–698.

Smith, F.M., B.D. Watts, J. Lyons, T. Keyes, B. Winn, A. Smith, F. Sanders & J. Thibault. 2019. Investigating Red Knot Migration Ecology along the Georgia and South Carolina Coasts: Spring 2019 Season Summaries. Center for Conservation Biology Technical Report Series: CCBTR-20-02. College of William and Mary/Virginia Commonwealth University, Williamsburg, VA. 53 pp.

South Carolina Department of Natural Resources. 2015. South Carolina’s State Wildlife Action Plan. Columbia, South Carolina. https://www.dnr.sc.gov/swap/index.html

South Carolina Department of Natural Resources. 2018. Shorebird research underscores importance of South Carolina beaches. South Carolina Department of Natural Resources. https://www.dnr.sc.gov/news/2018/jun/jun7_shorebirds.html.

Studds, C.E., B.E. Kendall, N.J. Murray, H.B. Wilson, D.I. Rogers, R.S. Clemens, K. Gosbell, C.J. Hassell, R. Jessop, D.S. Melville, D.A. Milton, C.D.T. Minton, H.P. Possingham, A.C. Riegen, P. Straw, E.J. Woehler & R.A. Fuller. 2017. Rapid population decline in migratory shorebirds relying on Yellow Sea tidal mudflats as stopover sites. Nature Communications 8: 14895.

Sullivan, B.L., C.L. Wood, M.J. Iliff, R.E. Bonney, D. Fink & S. Kelling. 2009. eBird: a citizen-based bird observation network in the biological sciences. Biological Conservation 142: 2282–2292.

Takahashi, F., F.J. Sanders & P.G.R. Jodice. 2021. Spatial and temporal overlap between foraging shorebirds and spawning horseshoe crabs (*Limulus polyphemus*) in the Cape Romain-Santee Delta Region of the U.S. Atlantic Coast. Wilson Journal of Ornithology 133: 58–72.

Thibault, J. 2013. Assessing the status and use of Red Knots in South Carolina. USFWS Traditional Section 6 Grant Report. South Carolina Department of Natural Resources.

Thibault, J. & M. Levisen. 2013. Red Knot prey availability project report. South Carolina Department of Natural Resources, Marine Resources Research Institute, Charleston, SC. 15pp.

Thomas, K., R.G. Kvitek & C. Bretz. 2003. Effects of human activity on the foraging behavior of sanderlings *Calidris alba*. Conservation Biology 109: 67–71.

Tucker, A., C. McGowan, J. Lyons, A. DeRose-Wilson & L. Clark. 2021. Species‐specific demographic and behavioral responses to food availability during migratory stopover. Population Ecology 64: 19–34

Tuma, M.E. & A.N. Powell. 2021. The Southeastern U.S. as a complex of use sites for nonbreeding rufa Red Knots: fifteen years of band-encounter data. Wader Study 128: 265–273.

U.S. Fish and Wildlife Service. 2001. Endangered and Threatened Wildlife and Plants; Final Determination of Critical Habitat for Wintering Piping Plovers. 66 Fed. Reg. 36038–36143.

U.S. Fish and Wildlife Service. 2014. Endangered and Threatened Wildlife and Plants; Threatened Species Status for the Rufa Red Knot. 79 Fed. Reg. 73706–73748.

van Gils, J.A., P.F. Battley, T. Piersma & R. Drent. 2005. Reinterpretation of gizzard sizes of Red Knots world-wide emphasises overriding importance of prey quality at migratory stopover sites. Proceedings of the Royal Society B 272: 2609–2618.

von Holle, B., J.L Irish, A. Spivy, J.F. Weishampel, A. Meylan, M.H. Godfrey, M. Dodd, S.H. Schweitzer, T. Keyes, F. Sanders, M.K. Chaplin & N.R. Taylor. 2019. Effects of future sea level rise on coastal habitat. Journal of Wildlife Management 83: 694–704.

White, G.C. & K.P. Burnham. 1999. Program MARK: survival estimation from populations of marked animals. Bird Study 46: S120–139.

Wilcove, D.S. & M. Wikelski. 2008. Going, Going, Gone: Is Animal Migration Disappearing. PLoS Biology 6: e188.

Xu, Y., Y. Si, J. Takekawa, Q. Liu, H.H.T. Prins, S. Yin, D.J. Prosser, P. Gong & W.F. de Boer. 2019. A network approach to prioritize conservation efforts for migratory birds. Conservation Biology 34: 416–426.

Verkuil, Y.I., E. Tavares, P.M. González, K. Choffe, O. Haddrath, M. Peck, L.J. Niles, A.J. Baker, T. Piersma & J.R. Conklin. 2021. Genetic structure in the nonbreeding range of *rufa* Red Knots suggests distinct Arctic breeding populations. Ornithological Applications 124: duab053.

